# Jumping back and forth: anthropozoonotic and zoonotic transmission of SARS-CoV-2 on mink farms

**DOI:** 10.1101/2020.09.01.277152

**Authors:** Bas B. Oude Munnink, Reina S. Sikkema, David F. Nieuwenhuijse, Robert Jan Molenaar, Emmanuelle Munger, Richard Molenkamp, Arco van der Spek, Paulin Tolsma, Ariene Rietveld, Miranda Brouwer, Noortje Bouwmeester-Vincken, Frank Harders, Renate Hakze-van der Honing, Marjolein C.A. Wegdam-Blans, Ruth J. Bouwstra, Corine GeurtsvanKessel, Annemiek A. van der Eijk, Francisca C. Velkers, Lidwien A.M. Smit, Arjan Stegeman, Wim H.M. van der Poel, Marion P.G. Koopmans

## Abstract

The zoonotic origin of the SARS-CoV-2 pandemic is still unknown. Animal experiments have shown that non-human primates, cats, ferrets, hamsters, rabbits and bats can be infected by SARS-CoV-2. In addition, SARS-CoV-2 RNA has been detected in felids, mink and dogs in the field. Here, we describe an in-depth investigation of outbreaks on 16 mink farms and humans living or working on these farms, using whole genome sequencing. We conclude that the virus was initially introduced from humans and has evolved, most likely reflecting widespread circulation among mink in the beginning of the infection period several weeks prior to detection. At the moment, despite enhanced biosecurity, early warning surveillance and immediate culling of infected farms, there is ongoing transmission between mink farms with three big transmission clusters with unknown modes of transmission. We also describe the first animal to human transmissions of SARS-CoV-2 in mink farms.

**One sentence summary:** SARS-CoV-2 transmission on mink farms.

## Main text

Late December 2019, SARS-CoV-2 was identified as the causative agent in a viral pneumonia outbreak, possibly related to a seafood and a live animal market in Wuhan, China (*1*). Since then, SARS-CoV-2 spread across the world and by August 31^rd^, over 25,200,000 people had been infected with SARS-CoV-2 resulting in over 840,000 deaths (*2*). In the Netherlands, over 72,000 infections have been confirmed, over 6,200 SARS-CoV-2 related deaths have been reported, and drastic measures have been put into place to prevent further spread of SARS-CoV-2 (*3*).

In view of the similarities with SARS-CoV-1, a zoonotic origin of the outbreak was suspected by the possible link with the Wuhan market where various animals were sold including fish, shellfish, poultry, wild birds and exotic animals. The finding of cases with onset of illness well before the period observed in the cluster, however, suggests the possibility of other sources (*4*). Although closely related coronaviruses in bats (*5, 6*) and pangolins (*7, 8*) have most sequence identity to SARS-CoV-2, the most likely diversion date of SARS-CoV-2 from the most closely related bat sequence is estimated to date back to somewhere between 1948-1982 (*9*). Therefore, the animal reservoir(s) of SARS-CoV-2 is (are) yet to be identified.

Experimental infections in dogs (*10*), cats (*10, 11*), ferrets (*10, 12*), hamsters (*13, 14*), rhesus macaques (*15*), tree shrew (*16*), cynomolgus macaques (*17*), grivets (*18*), common marmosets (*19*), rabbits (*20*), and fruit bats (*21*) have shown that these species are susceptible to SARS-CoV-2, and experimentally infected cats, tree shrews, hamsters and ferrets could transmit the virus. In contrast, experimental infection of pigs and several poultry species with SARS-CoV-2 proved to be unsuccessful (*10, 21, 22*). SARS-CoV-2 has also sporadically been identified in naturally infected animals. In the USA and in Hong Kong, SARS-CoV-2 RNA has been detected in dogs (*23*). In the Netherlands, France, Hong Kong, Belgium and the USA, cats have tested positive by RT-PCR for SARS-CoV-2 (*24*–*27*). Furthermore, SARS-CoV-2 has been detected in four tigers and three lions in a zoo in New York (*28*). In Italy, the Netherlands and in Wuhan, antibodies to SARS-CoV-2 have been detected in cats (*29*–*31*). Recently, detected SARS-CoV-2 was detected in farmed mink (*Neovison vison*) that showed signs of respiratory disease and increased mortality (*29, 32*).

Thereafter, the Dutch national response system for zoonotic diseases was activated, and it was concluded that the public health risk of animal infection with SARS-CoV-2 was low, but that there was a need for increased awareness of possible involvement of animals in the COVID-19 epidemic. Therefore, from May 20^th^ 2020 onwards, mink farmers, veterinarians and laboratories were obliged to report symptoms in mink (family *Mustelidae*) to the Netherlands Food and Consumer Product Safety Authority (NFCPSA) and an extensive surveillance system was set up (*33*).

Whole genome sequencing (WGS) can be used to monitor the emergence and spread of pathogens (*34*–*37*). As part of the surveillance effort in the Netherlands over 1,750 SARS-CoV-2 viruses have been sequenced to date from patients from different parts of the Netherlands (*38*). Here, we describe an in-depth investigation into the SARS-CoV-2 outbreak in mink farms and mink farm employees in the Netherlands, combining epidemiological information, surveillance data and WGS on the human-animal interface.

## Methods

### Outbreak investigation

Following initial detection of SARS-CoV-2 in mink on two farms on April 23^rd^ and April 25^th^, respectively, as part of routine health monitoring done by the Royal GD Animal Health service and subsequent investigation by Wageningen Bioveterinary Research (WBVR), the national reference laboratory for notifiable animal diseases, a One Health outbreak investigation team was convened (*39, 40*). Subsequently, respiratory signs and increased mortality in mink was made notifiable by the Dutch Ministry of Agriculture, Nature and Food Quality and the farms were quarantined (no movements of animals and manure and visitor restrictions). On May 7^th^ two other mink farms in the same region were confirmed to be infected. By the end of May the Dutch minister of Agriculture decided that all mink on SARS-CoV-2 infected farms had to be culled. Moreover, as the clinical manifestation of the infection was highly variable within and between farms, including asymptomatic infections, weekly testing of dead animals for SARS-CoV-2 infections became compulsory for all mink farms in the Netherlands. Moreover, a nation-wide transport ban of mink and mink manure, and a strict hygiene and visitor protocol was implemented. The first infected mink farms were culled from June 6^th^ onwards. From the 10^th^ infected farm (NB10) onwards, culling took place within 1-3 days after diagnosis. In this manuscript, the data up to June 26^th^, when a total of 16 mink farms in the Netherlands were found positive for SARS-CoV-2 infections, is presented.

### Veterinary and human contact tracing

The Netherlands Food and Consumer Product Safety Authority (NVWA) traced animal related contacts with other mink farms. Backward and forward tracing of possible high-risk contacts was done in the framework of the standard epidemiological investigation by the NVWA (i.e. focused on movement of vehicles, visitors such as veterinary practitioners, (temporary) workers, sharing of equipment between farms and transport and delivery of materials, such as feed, pelts, carcasses and manure). Persons with possible exposure from this investigation, as well as farm owners and resident farm workers were asked to report health complaints to the municipal health service for testing and – in the case of confirmed infections – for health advice and further contact tracing. Farm owners and workers on infected mink farms were informed of potential risks and were given advice on the importance and use of personal protective equipment and hygiene when handling animals (*41*). The contact structure on the farms was assessed through in-depth interviews, to identify additional persons with possible exposure to mink. In order to provide an enhanced set of reference genome sequences, anonymized samples from patients that had been diagnosed with COVID-19 in the area of the same four-digits postal codes as farms NB1-NB4 in March and April 2020 were retrieved from clinical laboratories in the region.

### SARS-CoV-2 diagnostics and sequencing

The presence of viral RNA in mink samples was determined using a RT-PCR targeting the E gene as previously described (*42*). For the human samples, diagnostic RT-PCR was performed for the E and the RdRp gene (*42*). In addition, serology was performed, using the Wantai Ig total and IgM ELISA, following the manufacturer’s instructions(*43*). For all samples with a Ct value <32, sequencing was performed using a SARS-CoV-2 specific multiplex PCR for Nanopore sequencing, as previously described (*3*). The libraries were generated using the native barcode kits from Nanopore (EXP-NBD104 and EXP-NBD114 and SQK-LSK109) and sequenced on a R9.4 flow cell multiplexing 24 samples per sequence run. Flow cells were washed and reused until less than 800 pores were active. The resulting raw sequence data was demultiplexed using Porechop (https://github.com/rrwick/Porechop). Primers were trimmed after which a reference-based alignment was performed. The consensus genome was extracted and positions with a coverage <30 were replaced with an “N” as described previously (*44*). Mutations in the genome compared to the GISAID sequence EPI_ISL_412973 were confirmed by manually checking the mapped reads and homopolymeric regions were manually checked and resolved by consulting reference genomes. The average SNP difference was determined using snp-dists (https://github.com/tseemann/snp-dists). All sequences generated in this study are available on GISAID.

### Phylogenetic analysis

All available near full-length Dutch SARS-CoV-2 genomes available on 1^st^ of July were selected (n=1,775) and aligned with the sequences from this study using MUSCLE (*45*). Sequences with >10% “Ns” were excluded. The alignment was manually checked for discrepancies after which IQ-TREE (*46*) was used to perform a maximum likelihood phylogenetic analysis under the GTR+F+I +G4 model as best predicted model using the ultrafast bootstrap option with 1,000 replicates. The phylogenetic trees were visualized in Figtree (http://tree.bio.ed.ac.uk/software/figtree/). For clarity reasons all bootstrap values below 80 were removed. To look at potential relationships with migrant workers, also all Polish sequences from GISAID (*47*) were included in the alignment (**Supplementary table 1**).

**Table 1.** Overview of human sampling on SARS-CoV-2 positive mink farms.

| Farm: | First diagnosis in animals: | Date(s) of sampling employees and family members: | PCR positive (%) | Serology positive (%) | Employees and family members tested positive (PCR and/or serology) |
| --- | --- | --- | --- | --- | --- |
| NB1 | 24-04-2020 | 28-04-2020 – 11-05-2020 | 5/6 (83%) | 5/5 (100%) | 6/6 (100%) |
| NB2 | 25-04-2020 | 31-03-2020 – 30-04-2020 | 1/2 (50%) | 8/8 (100%) | 8/8 (100%) |
| NB3 | 07-05-2020 | 11-05-2020 – 26-05-2020 | 5/7 (71%) | 0/6 (0%)* | 5/7 (71%) |
| NB4 | 07-05-2020 | 08-05-2020 | 1/3 (33%) | 2/2 (100%) | 2/3 (66%) |
| NB5 | 31-05-2020 | 01-06-2020 | 2/7 (29%) | 3/6 (50%) | 3/7 (43%) |
| NB6 | 31-05-2020 | 01-06-2020 | 1/6 (17%) | 4/6 (66%) | 4/6 (66%) |
| **NB7** | 31-05-2020 | 10-06-2020 – 01-07-2020 | 8/10 (80%) | NA** | 8/10 (80%) |
| **NB8** | 02-06-2020 | 03-06-2020 | 5/10 (50%) | 5/9 (56%) | 8/10 (80%) |
| **NB9** | 04-06-2020 | 07-06-2020 | 1/7 (14%) | 1/7 (14%) | 2/7 (29%) |
| **NB10** | 08-06-2020 | 11-06-2020 | 1/8 (13%) | 3/8 (38%) | 4/8 (50%) |
| **NB11** | 08-06-2020 | 11-06-2020 | 1/3 (33%) | 0/2 (0%) | 1/3 (33%) |
| **NB12** | 09-06-2020 | 11-06-2020 | 6/9 (66%) | 2/8 (25%) | 7/9 (78%) |
| **NB13** | 14-06-2020 | 11-06-2020 – 18-06-2020 | 3/3 (33%) | 0/2 (0%) | 3/3 (33%) |
| **NB14** | 14-06-2020 | 14-06-2020 | 1/3 (100%) | 5/6 (83%) | 5/6 (83%) |
| **NB15** | 21-06-2020 | 10-06-2020 – 30-06-2020 | 2/2 (100%) | NA** | 2/2 (100%) |
| **NB16** | 21-06-2020 | 23-06-2020 | 0/2 (0%) | NA** | 0/2 (0%) |
| **Total:** |  |  | **43/88 (49%)** | **38/75 (51%)** | **66/97 (68%)** |

\* Serology was done approximately one week before the positive PCR test.
\*\* No serology was performed

### Mapping specific mutation patterns on mink farms and in mink farm employees

Amino acid coordinates are described in relation to the Genbank NC_045512.2 reference genome. Open reading frames were extracted from the genome alignment using the genome annotation as supplied with the reference genome. A custom R script was used to distinguish synonymous from non-synonymous mutations and non-synonymous mutations were visualized using a tile map from the ggplot2 package (*48*).

### Geographical overview of mink farms in the Netherlands and SARS-CoV-2 positive farms

To protect confidentiality, SARS-CoV-2 positive mink farms were aggregated at municipality level. The datasets “*Landbouw; gewassen, dieren en grondgebruik naar gemeente”* and “*Wijk-en Buurtkaart 2019”* from Statistics Netherlands (CBS) were used (*49*). Maps were created using R packages sp (*50*), raster (*51*) and rgdal (*52*) and ArcGIS 10.6 software by ESRI.

## Results

SARS-CoV-2 was first diagnosed on two mink farms in the Netherlands on April 23^rd^ (NB1) and April 25^th^ (NB2), respectively. After the initial detection of SARS-CoV-2 on these farms an in-depth investigation was started to look for potential transmission routes and to perform an environmental and occupational risk assessment. Here, we describe the results of the outbreak investigation of the first 16 SARS-CoV-2 infected mink farms by combining SARS-CoV-2 diagnostics, WGS and in-depth interviews.

### Screening of farm workers and contacts

Farm owners of the 16 SARS-CoV-2 positive mink farms were contacted by the municipal health services to conduct contact investigation and samples were taken for RT-PCR-based and serological SARS-CoV-2 diagnostics. In total, 97 individuals were tested by either serological assays and/or RT-PCR. In total, 43 out of 88 (49%) upper-respiratory tract samples tested positive by RT-PCR while 38 out of 75 (51%) serum samples tested positive for SARS-CoV-2 specific antibodies. In total, 66 of 97 (67%) of the persons tested had evidence for SARS-CoV-2 infection (**table 1**).

### Anthropozoonotic transmission of SARS-CoV-2

During the interview on April 28^th^, four out of five employees from NB1 reported that they had experienced respiratory symptoms before the outbreak was detected in minks, but none of them had been tested for SARS-CoV-2. The first day of symptoms of people working on NB1 ranged from April 1^st^ to May 9^th^. For 16 of the mink, sampled on April 28^th^, and one farm employee, sampled on May 4^th^, a WGS was obtained (hCov-19/Netherlands/NoordBrabant_177/2020). The human sequence clusters within the mink sequences although it had 7 nucleotides difference with the closest mink sequence (**Figure 1** and cluster A in **figures 2 and 3**). On farm NB2, SARS-CoV-2 was diagnosed on April 25^th^. Retrospective analysis showed that one employee from NB2 had been hospitalized with SARS-CoV-2 on March 31^st^. All samples from the 8 employees taken on April 30^th^ were negative by RT-PCR but tested positive for SARS-CoV-2 antibodies. The virus sequence obtained from animals was distinct from that of farm NB1, indicating a separate introduction (**Figure 2 and 3**, cluster B).

**Figure 1:**
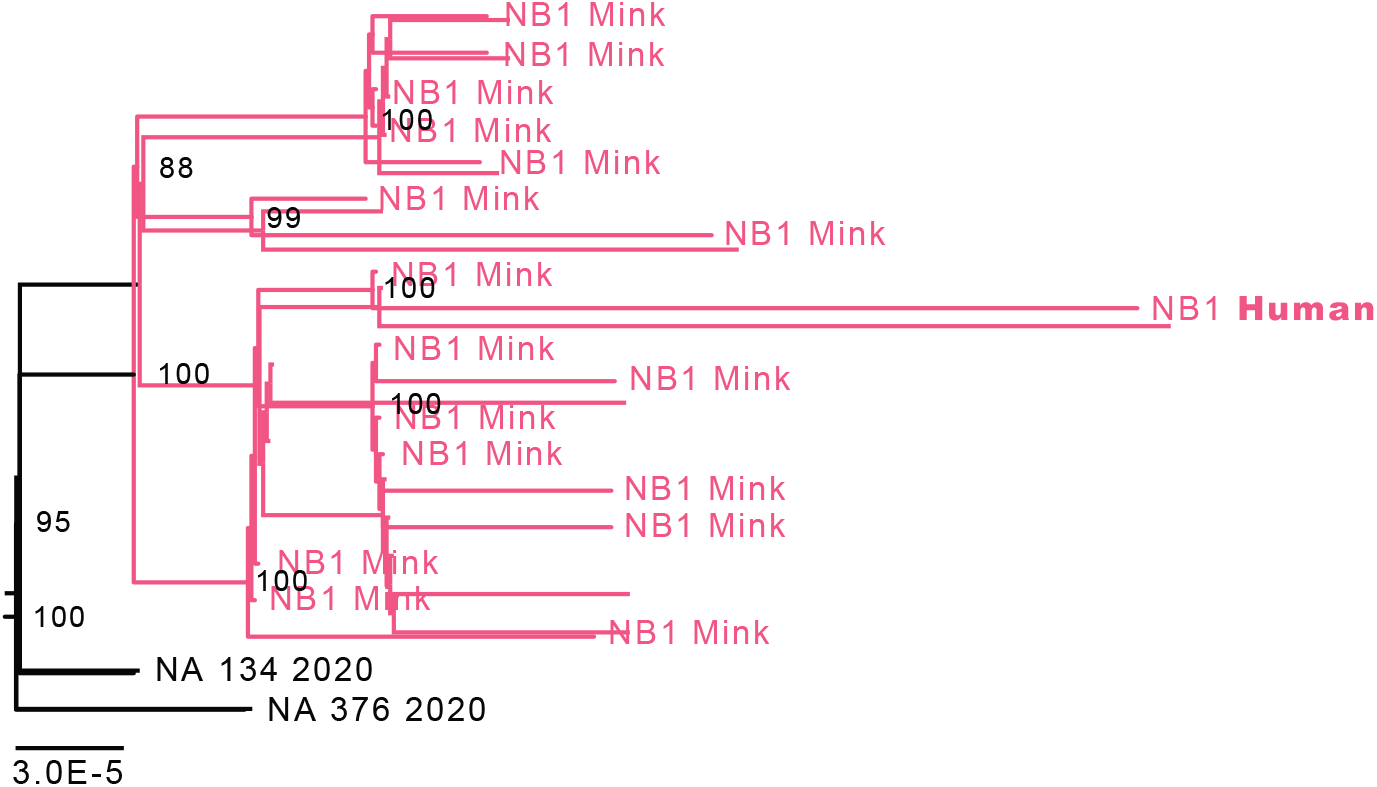
Zoom of the phylogenetic analysis of NB1. A maximum likelihood analysis was performed using all available SARS-CoV-2 Dutch sequences. Sequences from mink on NB1 are depicted in red and from the employee on NB1 in blue. The two sequences in black at the root of the cluster are the closest matching human genome sequences from the national SARS-CoV-2 sequence database. Scale bar represents units of substitutions per site.

**Figure 2:**
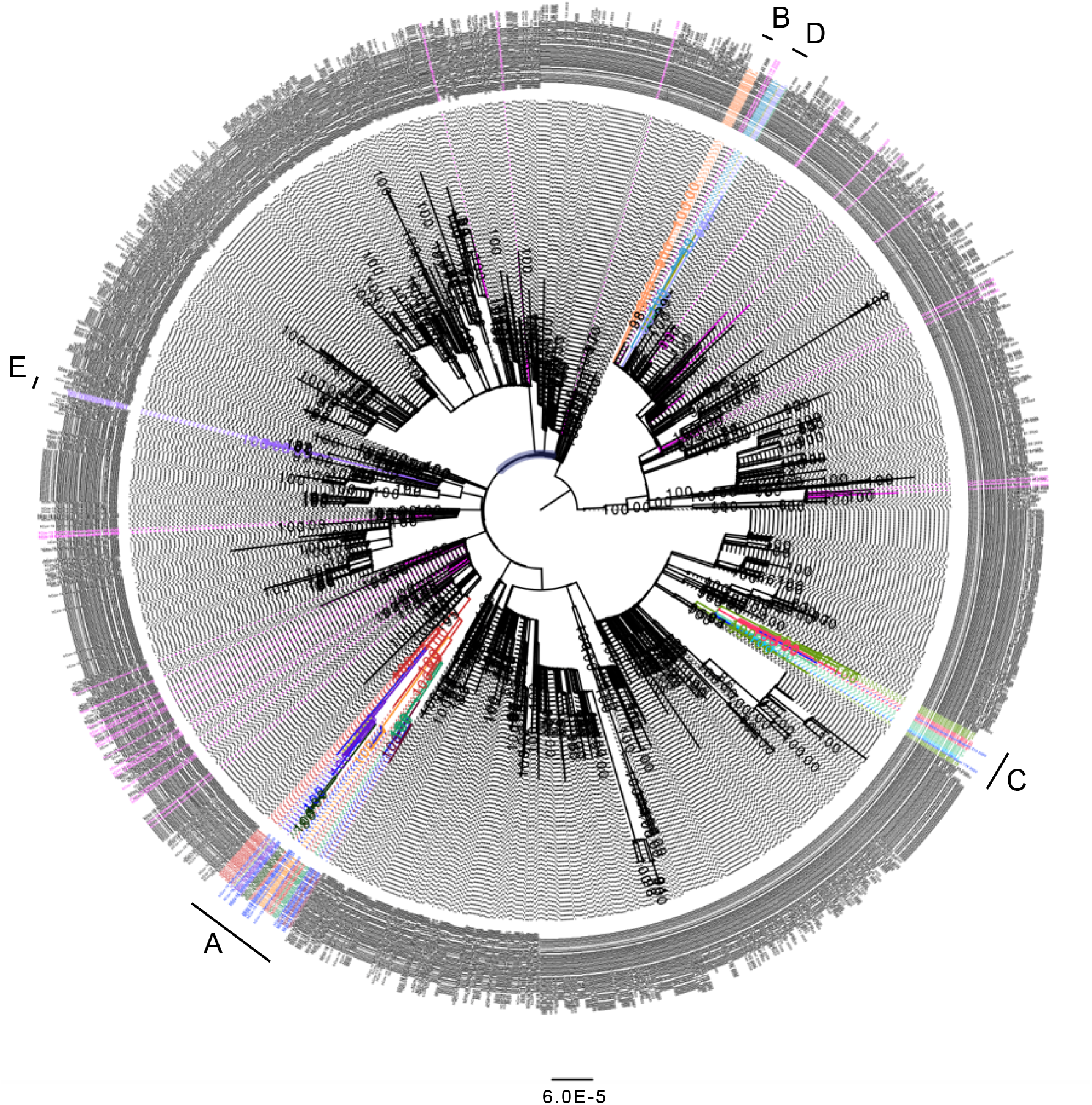
Maximum likelihood analysis of all SARS-CoV-2 Dutch sequences. The sequences derived from minks from different farms are indicated with different colors, human sequences related to the mink farms in blue and samples from similar 4-digit postal code are indicated in magenta. Scale bar represents units of substitutions per site.

**Figure 3:**
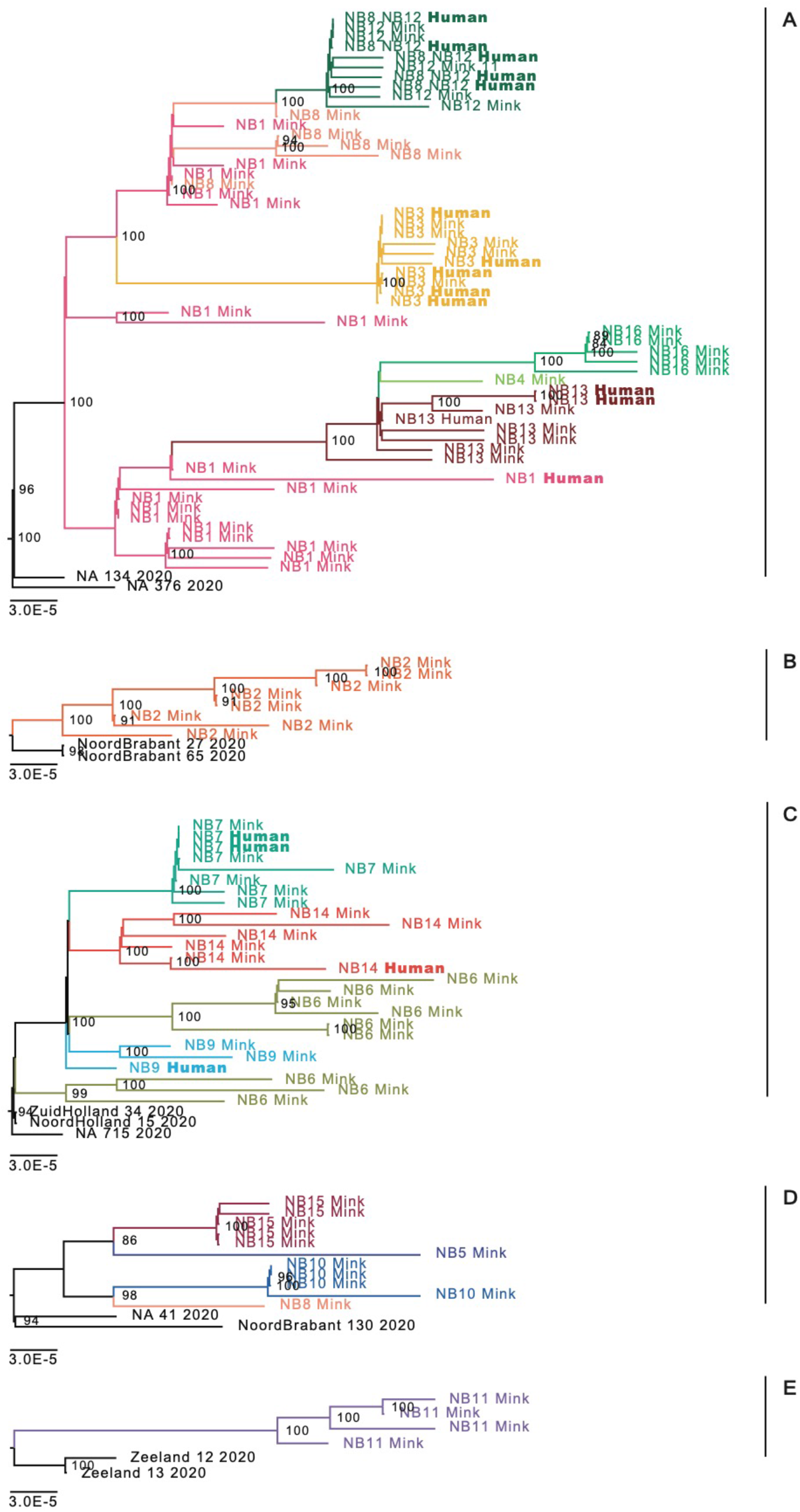
Phylogenetic analysis of SARS-CoV-2 strains detected in the 5 mink farm clusters. The sequences derived from different farms are depicted in different colors. Scale bar represents units of substitutions per site.

### Zoonotic transmission of SARS-CoV-2

On mink farm NB3 SARS-CoV-2 infection was diagnosed on May 7^th^. Initially all seven employees tested negative for SARS-CoV-2, but when retested between May 19^th^ and May 26^th^ after developing COVID-19 related symptoms, 5 out of 7 individuals working or living on the farm tested positive for SARS-CoV-2 RNA. WGS were obtained from these five individuals and the clustering of these sequences with the sequences derived from mink from NB3, together with initial negative test result and the start of the symptoms, indicate that the employees were infected with SARS-CoV-2 after the mink on the farm got infected. Also, an additional infection was observed based on contact-tracing: a close contact of one of the employees – who did not visit the farm – got infected with the SARS-CoV-2 strain found on NB3. Animal and human sequences from farm NB3 were related to those from farm NB1, but were both part of cluster A.

Similarly, on mink farm NB7 zoonotic transmission from mink to human most likely occurred. On this farm, SARS-CoV-2 infection in mink was diagnosed on May 31^st^ and employees initially tested negative for SARS-CoV-2 but started to develop symptoms at a later stage. Samples were taken between June 10^th^ and July 1^st^ from 10 employees of which 8 tested positive for SARS-CoV-2 RNA. From 2 samples WGS could be generated from the employees which clustered together with the sequences from the animals from this farm.

### Comparison with national reference database and enhanced regional sampling

The sequences generated from mink farms and from mink farm employees were compared with the national database consisting of around 1,775 WGS. In addition, to discriminate between locally acquired infections and mink farm related SARS-CoV-2 infection, and to determine the potential risk for people living close to mink farms, WGS was also performed on 34 SARS-CoV-2 positive samples from individuals who live in the same four-digit postal code area compared to the first four mink farms. These local sequences reflected the general diversity seen in the Netherlands and were not related to the clusters of mink sequences found on the mink farms, thereby also giving no indication of spill-over to people living in close proximity to mink farms (sequences shown in magenta, **Figure 2**). The sequences from the mink farm investigation were also compared to sequences from Poland (n=65), since many of the mink farm workers were seasonal migrants from Poland, but the these were not related.

### Mink farm related sequence clusters

Phylogenetic analysis of the mink SARS-CoV-2 genomes showed that mink sequences of 16 farms grouped into 5 different clusters (**Figure 2 and 3**). Viruses from farms NB1, NB3, NB4, NB8, NB12, NB13 and NB16 belonged to cluster A, sequences from NB2 were a separate cluster (B), those from farms NB6, NB7, NB9 and NB14 grouped together in cluster C, NB5, NB8, NB10 and NB15 grouped to cluster D, and NB11 had sequences designated as cluster E. On farm NB8, SARS-CoV-2 viruses could be found from both cluster A and cluster D. A detailed inventory of possible common characteristics, like farm owner, shared personnel, feed supplier and veterinary service provider, was made. In some cases, a link was observed with the same owners of several farms, for instance for cluster A for NB1 and NB4, and for NB8 and NB12. Although NB7, NB11 and NB15 were also linked to the same owner, viruses from these farms belonged to cluster C, D and E respectively. No common factor could be identified for most farms and clustering could also not be explained by geographic distances as multiple clusters were detected in different farms located close to each other (**Table 2 and figure 4**).

**Table 2.** Overview of the clusters detected on the different farms.

| Farm: | Date of diagnosis: | Sequence cluster: | Same owner: | Feed supplier: | Vet**: | Number of sequences (human): | Sequence diversity (average): | Mink population size: | Detection***: |
| --- | --- | --- | --- | --- | --- | --- | --- | --- | --- |
| NB1 | 24-04-20 | A | NB1, NB4 | 1 | I | 17 (1) | 0-9 (3.9) | 75,711 | Notification |
| NB2 | 25-04-20 | B |  | 1 | II | 8 | 0-8 (3.6) | 50,473 | Notification |
| NB3 | 07-05-20 | A |  | 2 | III | 5 (5) | 0-2 (0.6) | 12,400 | Notification |
| NB4 | 07-05-20 | A | NB1, NB4 | 1 | I | 1 | NA | 67,945 | Contact tracing NB1 |
| NB5 | 31-05-20 | D |  | 1 | IV | 1 | NA | 38,936 | EWS-Ser+PM-1st |
| NB6 | 31-05-20 | C |  | 3 | V | 9 | 0-12 (6.8) | 54,515 | EWS-Ser+PM-1st |
| NB7 | 31-05-20 | C | NB7, NB11, NB15 | 3 | II | 6 (2) | 0-4 (1.4) | 79,355 | EWS-PM-1st |
| NB8 | 02-06-20 | A/D | NB8, NB12* | 3 | V | 6 (5) | 0-6 (2.6) | 39,144 | EWS-Ser+PM-1st |
| NB9 | 04-06-20 | C |  | 2 | V | 2 (1) | 0-3 (1.5) | 32,557 | EWS-Ser+PM-2nd |
| NB10 | 08-06-20 | D |  | 3 | II | 4 | 0-3 (1.1) | 26,824 | EWS-Ser+PM-2nd |
| **NB11** | 08-06-20 | E | NB7,<br>NB11,<br>NB15 | 3 | II | 4 | 0-4 (2.2) | 38,745 | EWS-PM-2nd |
| **NB12** | 09-06-20 | A | NB8,<br>NB12* | 3 | II | 5 | 0-3 (1.2) | 55,352 | Notification |
| **NB13** | 14-06-20 | A |  | 3 | V | 5 (3) | 0-5 (3.2) | 20,366 | EWS-PM-5th |
| **NB14** | 14-06-20 | C |  | 3 | II | 5 (1) | 0-7 (3.7) | 28,375 | EWS-PM-5th |
| **NB15** | 21-06-20 | D | NB7,<br>NB11,<br>NB15 | 3 | II | 5 | 0-2 (0.6) | 35,928 | EWS-PM-6th |
| **NB16** | 21-06-20 | A |  | 3 | II | 5 | 0-4 (1.6) | 66,920 | EWS-PM-6th |

\* There was exchange of personnel in these two locations.
\*\* Veterinarian II and V were both from the same veterinary practice.
\*\*\* Notification: based on reporting of clinical signs which was obligated from 26 April onwards; EWS-Ser-Detection based on a one-off nation-wide compulsory serological screening of all mink farms at the end of May/early June by GD Animal Health; EWS-PM-Detection based on the early warning monitoring system for which carcasses of animals that died of natural causes were submitted weekly for PCR testing by GD Animal Health from the end of May onwards in a weekly cycle (EWS-PM 1<sup>st</sup> to 6<sup>th</sup> post mortem screening).

**Figure 4.**
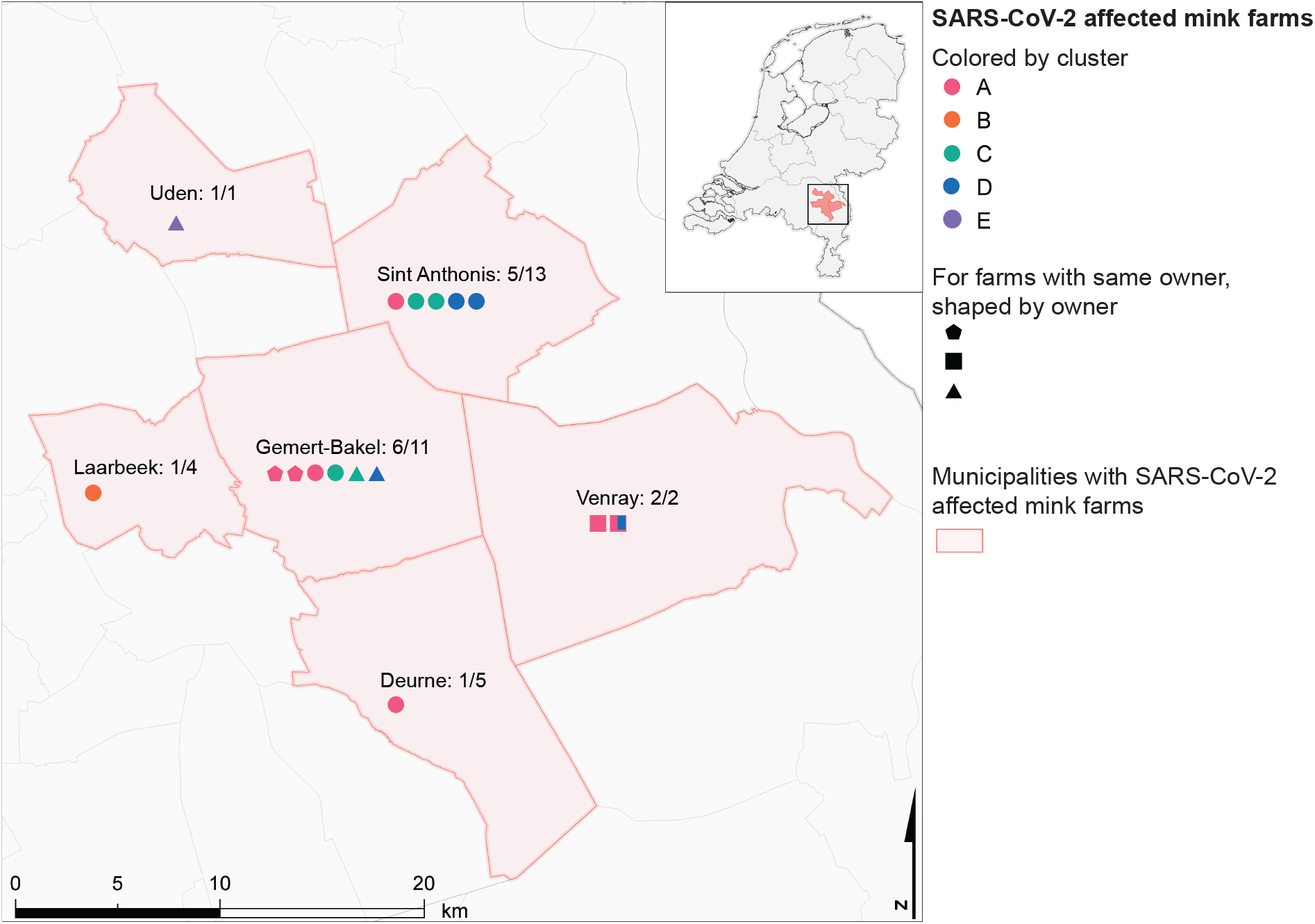
Geographical overview of SARS-CoV-2 positive mink farms per municipality affected. The proportion of SARS-CoV-2 positive mink farms over the total number of mink farms (CBS, 2019) is indicated. Symbols for positive farms are colored by cluster and shapes indicate farms with a same owner.

In total 18 sequences from mink farm employees or close contacts were generated from seven different farms. In most cases, these human sequences were near-identical to the mink sequences from the same farm. For NB1 the situation was different and the human sequence clusters deeply within the sequences derived from mink (Figure 1), with 7 nucleotides difference with the closest related mink sequence. This was also the case on farm NB14, with 4 nucleotides difference with the closest related mink sequence. Employees sampled at mink farm NB8 clustered with animals from NB12 which can be explained by the exchange of personnel between these two farms.

### Within farm diversity

SARS-CoV-2 was detected on mink farm NB1-NB4 after reports of respiratory symptoms and increased mortality in mink. The sequences from farm NB1 had between 0 and 9 nucleotides differences (average 3.9 nucleotides) and from NB2 between 0 and 8 nucleotides differences (average of 3.6), which is much more than what has been observed in outbreaks in human settings. The sequences from NB3 had 0 to 2 nucleotides difference suggesting that the virus was recently introduced, in line with the observed disease in humans, which occurred in the weeks post diagnosis of the infection in mink. After the initial detection of SARS-CoV-2 on mink farms, farms were screened weekly. The first, second, fifth and sixth weekly screening yielded new positives. The sequences of mink at NB6 had between 0 and 12 nucleotides differences, whereas diversity was lower for the subsequent farm sequences (Table 2).

Several non-synonymous mutations were identified among the mink sequences compared to the Wuhan reference sequence NC_045512.2. However, no particular amino acid substitutions were found in all mink samples (**Figure 5**). Of note, three of the clusters had the position 614G variant (clusters A, C and E), and 2 had the original variant. There were no obvious differences in the presentation of disease in animals or humans between clusters based on the data available at this stage, but further data collection and analysis, also for cases after NB16, are ongoing to investigate this further. The observed mutations can also be found in the general population and the same mutations also were found in human cases which were related to the mink farms.

**Figure 5.**
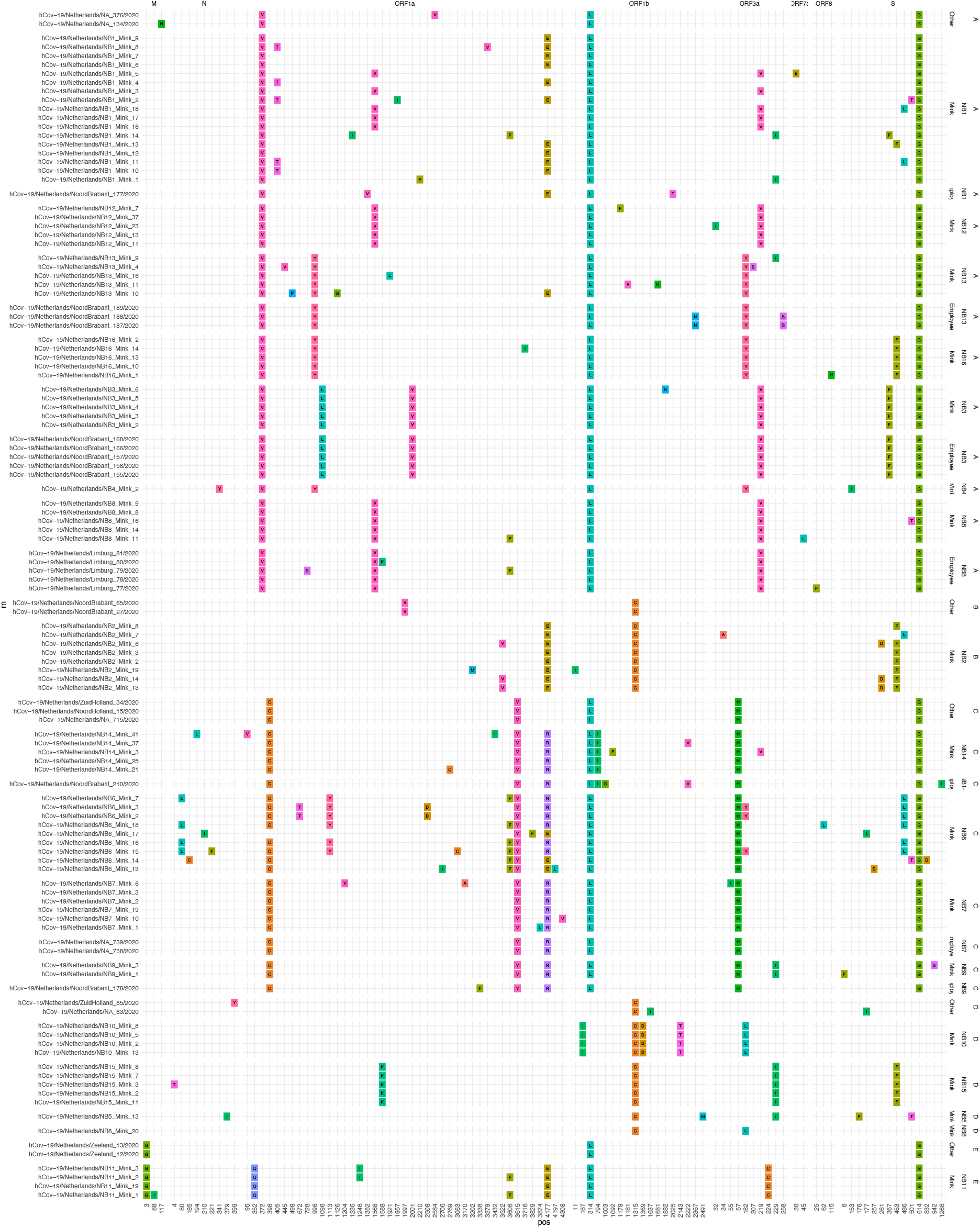
Overview of the specific amino acid mutations found in mink farms. Above the x-axis the open reading frames (ORF) are indicated and on the x-axis the amino acid position within each ORF is indicated. On the y-axis the sequence names are indicated and on the right side of the graph the cluster numbers and specific farm identifiers and the type of host are used to group the samples.

## Discussion

Here we show ongoing SARS-CoV-2 transmission in mink farms and spill-over events to humans. To the best of our knowledge, these are the first animal to human SARS-CoV-2 transmission events documented. More research in minks and other mustelid species, to demonstrate if these species can be a true reservoir of SARS-CoV-2 although from our observations we consider this likely. After the detection of SARS-CoV-2 on mink farms, 68% of the tested farm workers and/or relatives or contacts were shown to be infected with SARS-CoV-2, indicating that contact with SARS-CoV-2 infected mink is a risk factor for contracting COVID-19.

A high diversity in the sequences from some mink farms was observed which most likely can be explained by many generations of infected animals before an increase in mortality was observed. The current estimates are that the substitution rate of SARS-CoV-2 is around 1.16*10^-3 substitutions/site/year (*53*), which corresponds to around one mutation per two weeks. This could mean that the virus was already circulating in mink farms for some time. However, there was also a relatively high sequence diversity observed in farms which still tested negative one week prior, hinting towards a faster evolution of the virus in the mink population. This can indicate that the virus might replicate more efficiently in mink or might have acquired mutations which makes the virus more virulent. However, no specific mutations were found in all mink samples, making increased virulence less likely. In addition, mink farms have large populations of animals which could lead to very efficient virus transmission. Generation intervals for SARS-CoV-2 in humans have been estimated to be around 4-5 days(*54*), but with high dose exposure in a high-density farm could potentially be shorter. Recently, a specific mutation in the spike protein (D614G) was shown to result in an increased virulence *in vitro* (*55*), while it was not associated an increased growth rate for cluster nor an increased mortality (*56*). This mutation was present in farm clusters A, C and E, but no obvious differences in clinical presentation, disease severity, or rate of transmission to humans was observed.

While we found sequences matching with the animal sequences on several farms, not all of these can be considered direct zoonotic transmissions. For instance, the two employees from mink farm NB3 were most likely infected while working at the mink farm given the specific clustering in the phylogenetic tree and the timing of infection. Subsequent human infections may have originated from additional zoonotic infections, or from human to human transmission within their household. Further proof that animals were the most likely source of infection was provided by the clear phylogenetic separation between farm related human cases and animal cases, from sequences from cases within the same 4-digit postal code area. Spill-back into the community living in the same 4-digit postal code area was not observed using sequence data, but cannot be entirely ruled out as the testing strategy during April and May was focusing on health care workers, persons with more severe symptoms, and persons at risk for complications, rather than monitoring community transmission and milder cases.

While the number of SARS-CoV-2 infected individuals was decreasing in the Netherlands in May and June, an increase in detection of SARS-CoV-2 in mink farms was observed. Based on WGS these sequences are part of multiple individual transmission chains linked to the mink farms and are not a reflection of the situation in the human population during this time. In some cases, the farms had the same owner but in other cases no epidemiological link could be established. People coming to the different farms might be a source but also semi-wild cats roaming around the farms or wildlife might play a role (*27*). So far, the investigation failed to identify common factors that might explain farm to farm spread. During interviews, it became clear that farms had occasionally hired temporary workers that had not been included in the testing and were lost to follow-up, stressing the need for vigorous biosecurity and occupational health guidance. Since our observation, SARS-CoV-2 infections have also been described in mink farms in Denmark, Spain and the USA (*57*–*59*), and mink farming is common in other regions of the world as well, also in China where around 26 million mink pelts are produced on a yearly basis (*60*). The population size and the structure of mink farms is such that it is conceivable that SARS-CoV-2 – once introduced – could continue to circulate. Therefore, continued monitoring and cooperation between human and animal health services is crucial to prevent the animals serving as a reservoir for continued infection in humans.

## Authors contributions

Conceptualization: B.B.O.M, R.S.S, A.S., M.P.G.K; investigation: B.B.O.M, D.F.N., R.S.S, A.S., M.P.G.K, L.A.M.S, W.H.N.v.d.P, R.J.M, R.J.B, E.M, R.M, A.v.d.S, P.T, A.R, M.B, N.B-V, F.H, R.H.v.d.H, M.C.A.W-B, R.J.B, C.G, A.A.v.d.E, F.C.V, L.A.M.S; supervision: M.P.G.M; writing original draft: B.B.O.M, R.S.S, M.P.G.K; writing review and editing, all authors

## Data and material availability

All data, code and materials used described in this manuscript are publicly available.

## Supplements

**Supplementary Figure 1:**
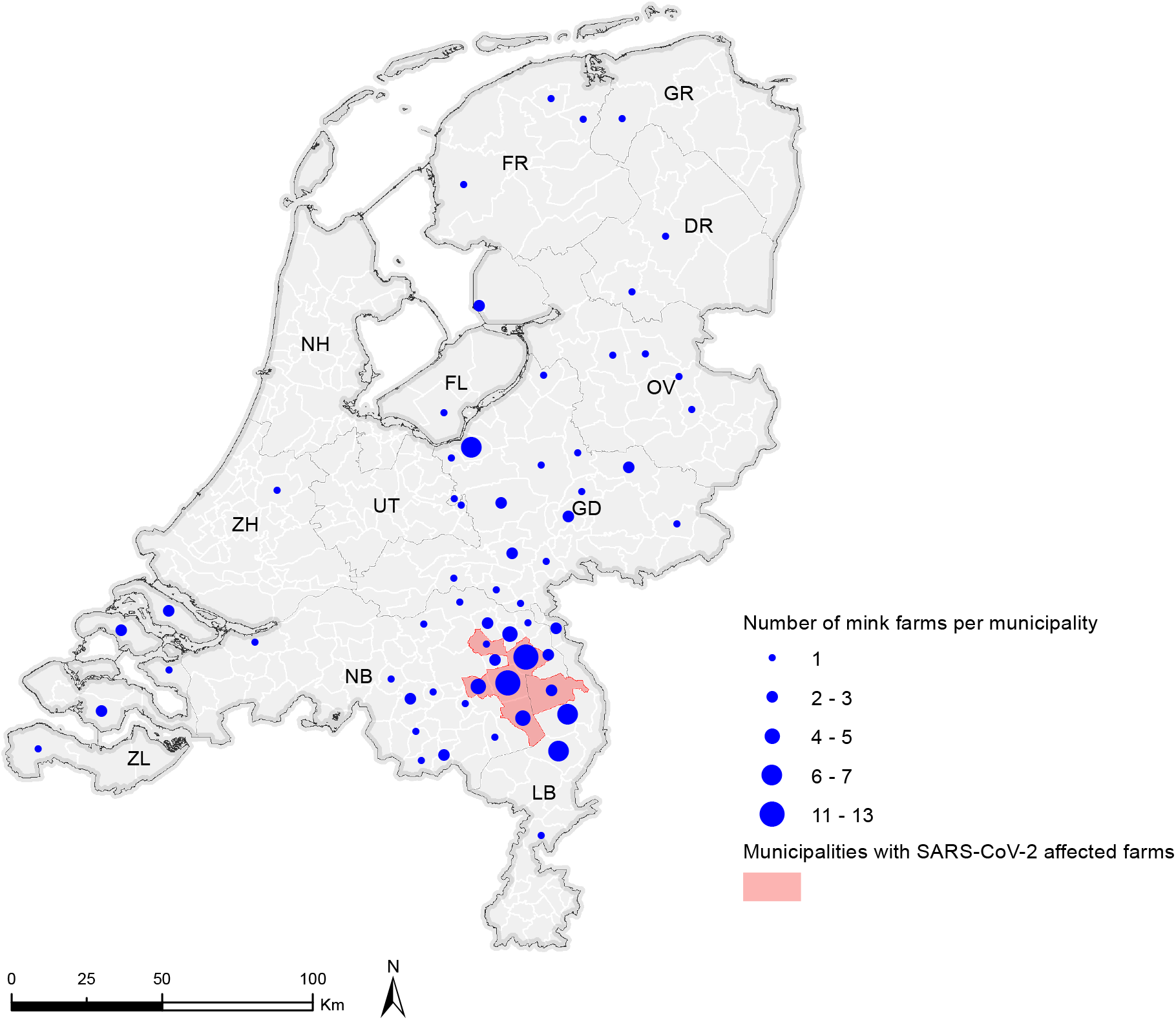
Number of mink farms per municipality in the Netherlands. Overview of the total number of mink farms per municipality (CBS, 2019). Municipalities with SARS-CoV-2 affected farms by June 21^st^ 2020 are shown in red.

**Supplementary table 1:** GISAID acknowledgement table.

## References

1. N. Zhu, D. Zhang, W. Wang, X. Li, B. Yang, J. Song, X. Zhao, B. Huang, W. Shi, R. Lu, P. Niu, F. Zhan, X. Ma, D. Wang, W. Xu, G. Wu, G. F. Gao, W. Tan, A novel coronavirus from patients with pneumonia in China, 2019. N. Engl. J. Med. 382, 727–733 (2020).

2. E. Dong, H. Du, L. Gardner, An interactive web-based dashboard to track COVID-19 in real time. Lancet Infect. Dis. 0 (2020),, doi: 10.1016/S1473-3099(20)30120-1.

3. B. B. Oude Munnink, D. F. Nieuwenhuijse, M. Stein, Á. O’Toole, M. Haverkate, M. Mollers, S. K. Kamga, C. Schapendonk, M. Pronk, P. Lexmond, A. van der Linden, T. Bestebroer, I. Chestakova, R. J. Overmars, S. van Nieuwkoop, R. Molenkamp, A. A. van der Eijk, C. GeurtsvanKessel, H. Vennema, A. Meijer, A. Rambaut, J. van Dissel, R. S. Sikkema, A. Timen, M. Koopmans, G. J. A. P. M. Oudehuis, J. Schinkel, J. Kluytmans, M. Kluytmans-van den Bergh, W. van den Bijllaardt, R. G. Berntvelsen, M. M. L. van Rijen, P. Schneeberger, S. Pas, B. M. Diederen, A. M. C. Bergmans, P. A. V. van der Eijk, J. Verweij, A. G. N. Buiting, R. Streefkerk, A. P. Aldenkamp, P. de Man, J. G. M. Koelemal, D. Ong, S. Paltansing, N. Veassen, J. Sleven, L. Bakker, H. Brockhoff, A. Rietveld, F. Slijkerman Megelink, J. Cohen Stuart, A. de Vries, W. van der Reijden, A. Ros, E. Lodder, E. Verspui-van der Eijk, I. Huijskens, E. M. Kraan, M. P. M. van der Linden, S. B. Debast, N. Al Naiemi, A. C. M. Kroes, M. Damen, S. Dinant, S. Lekkerkerk, O. Pontesilli, P. Smit, C. van Tienen, P. C. R. Godschalk, J. van Pelt, A. Ott, C. van der Weijden, H. Wertheim, J. Rahamat-Langendoen, J. Reimerink, R. Bodewes, E. Duizer, B. van der Veer, C. Reusken, S. Lutgens, P. Schneeberger, M. Hermans, P. Wever, A. Leenders, H. ter Waarbeek, C. Hoebe, Rapid SARS-CoV-2 whole-genome sequencing and analysis for informed public health decision-making in the Netherlands. Nat. Med., 1–6 (2020).

4. F. Wu, S. Zhao, B. Yu, Y. M. Chen, W. Wang, Z. G. Song, Y. Hu, Z. W. Tao, J. H. Tian, Y. Y. Pei, M. L. Yuan, Y. L. Zhang, F. H. Dai, Y. Liu, Q. M. Wang, J. J. Zheng, L. Xu, E. C. Holmes, Y. Z. Zhang, A new coronavirus associated with human respiratory disease in China. Nature. 579, 265–269 (2020).

5. H. Zhou, X. Chen, T. Hu, J. Li, H. Song, Y. Liu, P. Wang, D. Liu, J. Yang, E. C. Holmes, A. C. Hughes, Y. Bi, W. Shi, A Novel Bat Coronavirus Closely Related to SARS-CoV-2 Contains Natural Insertions at the S1/S2 Cleavage Site of the Spike Protein. Curr. Biol. (2020), doi: 10.1016/j.cub.2020.05.023.

6. P. Zhou, X. Lou Yang, X. G. Wang, B. Hu, L. Zhang, W. Zhang, H. R. Si, Y. Zhu, B. Li, C. L. Huang, H. D. Chen, J. Chen, Y. Luo, H. Guo, R. Di Jiang, M. Q. Liu, Y. Chen, X. R. Shen, X. Wang, X. S. Zheng, K. Zhao, Q. J. Chen, F. Deng, L. L. Liu, B. Yan, F. X. Zhan, Y. Y. Wang, G. F. Xiao, Z. L. Shi, A pneumonia outbreak associated with a new coronavirus of probable bat origin. Nature. 579, 270–273 (2020).

7. T. T. Y. Lam, M. H. H. Shum, H. C. Zhu, Y. G. Tong, X. B. Ni, Y. S. Liao, W. Wei, W. Y. M. Cheung, W. J. Li, L. F. Li, G. M. Leung, E. C. Holmes, Y. L. Hu, Y. Guan, Identifying SARS-CoV-2 related coronaviruses in Malayan pangolins. Nature, 1–6 (2020).

8. G. Z. Han, Pangolins Harbor SARS-CoV-2-Related Coronaviruses. Trends Microbiol. (2020),, doi: 10.1016/j.tim.2020.04.001.

9. M. F. Boni, P. Lemey, X. Jiang, T. T.-Y. Lam, B. Perry, T. Castoe, A. Rambaut, D. L. Robertson, bioRxiv, in press, doi: 10.1101/2020.03.30.015008.

10. J. Shi, Z. Wen, G. Zhong, H. Yang, C. Wang, B. Huang, R. Liu, X. He, L. Shuai, Z. Sun, Y. Zhao, P. Liu, L. Liang, P. Cui, J. Wang, X. Zhang, Y. Guan, W. Tan, G. Wu, H. Chen, Z. Bu, Susceptibility of ferrets, cats, dogs, and other domesticated animals to SARS– coronavirus 2. Science (80-.)., eabb7015 (2020).

11. P. J. Halfmann, M. Hatta, S. Chiba, T. Maemura, S. Fan, M. Takeda, N. Kinoshita, S.-I. Hattori, Y. Sakai-Tagawa, K. Iwatsuki-Horimoto, M. Imai, Y. Kawaoka, Transmission of SARS-CoV-2 in Domestic Cats. N. Engl. J. Med., NEJMc2013400 (2020).

12. M. Richard, A. Kok, D. de Meulder, T. M. Bestebroer, M. M. Lamers, N. M. A. Okba, M. F. van Vlissingen, B. Rockx, B. L. Haagmans, M. P. G. Koopmans, R. A. M. Fouchier, S. Herfst, bioRxiv, in press, doi: 10.1101/2020.04.16.044503.

13. S. F. Sia, L.-M. Yan, K. Fung, J. M. Nicholls, M. Peiris, H.-L. Yen, Pathogenesis and transmission of SARS-CoV-2 virus in golden Syrian hamsters SUBJECT AREAS Infectious Diseases Small Animal Medicine (2020), doi: 10.21203/rs.3.rs-20774/v1.

14. J. F. W. Chan, A. J. Zhang, S. Yuan, V. K. M. Poon, C. C. S. Chan, A. C. Y. Lee, W. M. Chan, Z. Fan, H. W. Tsoi, L. Wen, R. Liang, J. Cao, Y. Chen, K. Tang, C. Luo, J. P. Cai, K. H. Kok, H. Chu, K. H. Chan, S. Sridhar, Z. Chen, H. Chen, K. K. W. To, K. Y. Yuen, Simulation of the clinical and pathological manifestations of Coronavirus Disease 2019 (COVID-19) in golden Syrian hamster model: implications for disease pathogenesis and transmissibility. Clin. Infect. Dis. (2020), doi: 10.1093/cid/ciaa325.

15. V. J. Munster, F. Feldmann, B. N. Williamson, N. van Doremalen, L. Pérez-Pérez, J. Schulz, K. Meade-White, A. Okumura, J. Callison, B. Brumbaugh, V. A. Avanzato, R. Rosenke, P. W. Hanley, G. Saturday, D. Scott, E. R. Fischer, E. de Wit, Respiratory disease in rhesus macaques inoculated with SARS-CoV-2. Nature, 1–7 (2020).

16. Y. Zhao, J. Wang, D. Kuang, J. Xu, M. Yang, C. Ma, S. Zhao, J. Li, H. Long, K. Ding, J. Gao, J. Liu, H. Wang, H. Li, Y. Yang, W. Yu, J. Yang, Y. Zheng, D. Wu, S. Lu, H. Liu, X. Peng, bioRxiv, in press, doi: 10.1101/2020.04.30.029736.

17. B. Rockx, T. Kuiken, S. Herfst, T. Bestebroer, M. M. Lamers, B. B. Oude Munnink, D. de Meulder, G. van Amerongen, J. van den Brand, N. M. A. Okba, D. Schipper, P. van Run, L. Leijten, R. Sikkema, E. Verschoor, B. Verstrepen, W. Bogers, J. Langermans, C. Drosten, M. Fentener van Vlissingen, R. Fouchier, R. de Swart, M. Koopmans, B. L. Haagmans, Comparative pathogenesis of COVID-19, MERS, and SARS in a nonhuman primate model. Science (80-.)., eabb7314 (2020).

18. C. Woolsey, V. Borisevich, A. N. Prasad, K. N. Agans, D. J. Deer, N. S. Dobias, J. C. Heymann, S. L. Foster, C. B. Levine, L. Medina, K. Melody, J. B. Geisbert, K. A. Fenton, T. W. Geisbert, R. W. Cross, bioRxiv Prepr. Serv. Biol., in press, doi: 10.1101/2020.05.17.100289.

19. S. Lu, Y. Zhao, W. Yu, Y. Yang, J. Gao, J. Wang, D. Kuang, M. Yang, J. Yang, C. Ma, J. Xu, H. Li, S. Zhao, J. Li, H. Wang, H. Long, J. Zhou, F. Luo, K. Ding, D. Wu, Y. Zhang, Y. Dong, Y. Liu, Y. Zheng, X. Lin, L. Jiao, H. Zheng, Q. Dai, Q. Sun, Y. Hu, C. Ke, H. Liu, X. Peng, bioRxiv, in press, doi: 10.1101/2020.04.08.031807.

20. B. L. Haagmans, D. Noack, N. M. Okba, W. Li, C. Wang, R. de Vries, S. Herfst, D. de Meulder, P. van Run, B. Rijnders, C. Rokx, F. van Kuppeveld, F. Grosveld, C. GeurtsvanKessel, M. Koopmans, B. Jan Bosch, T. Kuiken, B. Rockx, bioRxiv, in press, doi: 10.1101/2020.08.24.264630.

21. K. Schlottau, M. Rissmann, A. Graaf, J. Schön, J. Sehl, C. Wylezich, D. Höper, T. C. Mettenleiter, A. Balkema-Buschmann, T. Harder, C. Grund, D. Hoffmann, A. Breithaupt, M. Beer, Experimental Transmission Studies of SARS-CoV-2 in Fruit Bats, Ferrets, Pigs and Chickens. SSRN Electron. J. (2020), doi: 10.2139/ssrn.3578792.

22. D. L. Suarez, M. J. Pantin-Jackwood, D. E. Swayne, S. A. Lee, S. M. Deblois, E. Spackman, bioRxiv, in press, doi: 10.1101/2020.06.16.154658.

23. T. H. C. Sit, C. J. Brackman, S. M. Ip, K. W. S. Tam, P. Y. T. Law, E. M. W. To, V. Y. T. Yu, L. D. Sims, D. N. C. Tsang, D. K. W. Chu, R. A. P. M. Perera, L. L. M. Poon, M. Peiris, Infection of dogs with SARS-CoV-2. Nature, 1–6 (2020).

24. C. Sailleau, M. Dumarest, J. Vanhomwegen, M. Delaplace, V. Caro, A. Kwasiborski, V. Hourdel, P. Chevaillier, A. Barbarino, L. Comtet, P. Pourquier, B. Klonjkowski, J. C. Manuguerra, S. Zientara, S. Le Poder, First detection and genome sequencing of SARS-CoV-2 in an infected cat in France. Transbound. Emerg. Dis. (2020), doi: 10.1111/tbed.13659.

25. A. Newman, D. Smith, R. R. Ghai, R. M. Wallace, M. K. Torchetti, C. Loiacono, L. S. Murrell, A. Carpenter, S. Moroff, J. A. Rooney, C. Barton Behravesh, First Reported Cases of SARS-CoV-2 Infection in Companion Animals - New York, March-April 2020. MMWR. Morb. Mortal. Wkly. Rep. 69, 710–713 (2020).

26. Promed Post – ProMED-mail, (available at https://promedmail.org/promed-post/?id=7314521).

27. N. Oreshkova, R.-J. Molenaar, S. Vreman, F. Harders, B. B. O. Munnink, R. Hakze, N. Gerhards, P. Tolsma, R. Bouwstra, R. Sikkema, M. Tacken, M. M. T. de Rooij, E. Weesendorp, M. Engelsma, C. Bruschke, L. A. M. Smit, M. Koopmans, W. H. M. van der Poel, A. Stegeman, bioRxiv, in press, doi: 10.1101/2020.05.18.101493.

28. R. Gollakner, I. Capua, Is COVID-19 the first pandemic that evolves into a panzootic? Vet. Ital. 56 (2020), doi: 10.12834/VetIt.2246.12523.1.

29. N. Oreshkova, R. J. Molenaar, S. Vreman, F. Harders, B. B. Oude Munnink, R. W. Hakze-van der Honing, N. Gerhards, P. Tolsma, R. Bouwstra, R. S. Sikkema, M. G. Tacken, M. M. de Rooij, E. Weesendorp, M. Y. Engelsma, C. J. Bruschke, L. A. Smit, M. Koopmans, W. H. van der Poel, A. Stegeman, SARS-CoV-2 infection in farmed minks, the Netherlands, April and May 2020. Eurosurveillance. 25, 2001005 (2020).

30. Q. Zhang, H. Zhang, K. Huang, Y. Yang, X. Hui, J. Gao, X. He, C. Li, W. Gong, Y. Zhang, C. Peng, X. Gao, H. Chen, Z. Zou, Z. Shi, M. Jin, bioRxiv, in press, doi: 10.1101/2020.04.01.021196.

31. E. I. Patterson, G. Elia, A. Grassi, A. Giordano, C. Desario, M. Medardo, S. L. Smith, E. R. Anderson, E. Lorusso, M. S. Lucente, G. Lanave, S. Lauzi, U. Bonfanti, A. Stranieri, V. Martella, fabrizio Solari Basano, V. R. Barrs, A. D. Radford, G. L. Hughes, S. Paltrinieri, N. Decaro, bioRxiv, in press, doi: 10.1101/2020.07.21.214346.

32. R. J. Molenaar, S. Vreman, R. W. Hakze-van der Honing, R. Zwart, J. de Rond, E. Weesendorp, L. A. M. Smit, M. Koopmans, R. Bouwstra, A. Stegeman, W. H. M. van der Poel, Clinical and Pathological Findings in SARS-CoV-2 Disease Outbreaks in Farmed Mink (Neovison vison). Vet. Pathol. (2020), doi: 10.1177/0300985820943535.

33. Bedrijfsmatig gehouden dieren en SARS-CoV-2 | Nieuws en media | NVWA, (available at https://www.nvwa.nl/nieuws-en-media/actuele-onderwerpen/corona/g/bedrijfsmatig-gehouden-dieren-en-corona).

34. B. B. Oude Munnink, E. Münger, D. F. Nieuwenhuijse, R. Kohl, A. van der Linden, C. M. E. Schapendonk, H. van der Jeugd, M. Kik, J. M. Rijks, C. B. E. M. Reusken, M. Koopmans, Genomic monitoring to understand the emergence and spread of Usutu virus in the Netherlands, 2016-2018. Sci. Rep. 10, 2798 (2020).

35. A. Arias, S. J. Watson, D. Asogun, E. A. Tobin, J. Lu, M. V. T. Phan, U. Jah, R. E. G. Wadoum, L. Meredith, L. Thorne, S. Caddy, A. Tarawalie, P. Langat, G. Dudas, N. R. Faria, S. Dellicour, A. Kamara, B. Kargbo, B. O. Kamara, S. Gevao, D. Cooper, M. Newport, P. Horby, J. Dunning, F. Sahr, T. Brooks, A. J. H. Simpson, E. Groppelli, G. Liu, N. Mulakken, K. Rhodes, J. Akpablie, Z. Yoti, M. Lamunu, E. Vitto, P. Otim, C. Owilli, I. Boateng, L. Okoror, E. Omomoh, J. Oyakhilome, R. Omiunu, I. Yemisis, D. Adomeh, S. Ehikhiametalor, P. Akhilomen, C. Aire, A. Kurth, N. Cook, J. Baumann, M. Gabriel, R. Wölfel, A. Di Caro, M. W. Carroll, S. Günther, J. Redd, D. Naidoo, O. G. Pybus, A. Rambaut, P. Kellam, I. Goodfellow, M. Cotten, Rapid outbreak sequencing of Ebola virus in Sierra Leone identifies transmission chains linked to sporadic cases. Virus Evol. 2, vew016 (2016).

36. N. R. Faria, M. U. G. Kraemer, S. C. Hill, J. G. de Jesus, R. S. Aguiar, F. C. M. Iani, J. Xavier, J. Quick, L. du Plessis, S. Dellicour, J. Thézé, R. D. O. Carvalho, G. Baele, C.-H. Wu, P. P. Silveira, M. B. Arruda, M. A. Pereira, G. C. Pereira, J. Lourenço, U. Obolski, L. Abade, T. I. Vasylyeva, M. Giovanetti, D. Yi, D. J. Weiss, G. R. W. Wint, F. M. Shearer, S. Funk, B. Nikolay, V. Fonseca, T. E. R. Adelino, M. A. A. Oliveira, M. V. F. Silva, L. Sacchetto, P. O. Figueiredo, I. M. Rezende, E. M. Mello, R. F. C. Said, D. A. Santos, M. L. Ferraz, M. G. Brito, L. F. Santana, M. T. Menezes, R. M. Brindeiro, A. Tanuri, F. C. P. dos Santos, M. S. Cunha, J. S. Nogueira, I. M. Rocco, A. C. da Costa, S. C. V. Komninakis, V. Azevedo, A. O. Chieppe, E. S. M. Araujo, M. C. L. Mendonça, C. C. dos Santos, C. D. dos Santos, A. M. Mares-Guia, R. M. R. Nogueira, P. C. Sequeira, R. G. Abreu, M. H. O. Garcia, A. L. Abreu, O. Okumoto, E. G. Kroon, C. F. C. de Albuquerque, K. Lewandowski, S. T. Pullan, M. Carroll, T. de Oliveira, E. C. Sabino, R. P. Souza, M. A. Suchard, P. Lemey, G. S. Trindade, B. P. Drumond, A. M. B. Filippis, N. J. Loman, S. Cauchemez, L. C. J. Alcantara, O. G. Pybus, Genomic and epidemiological monitoring of yellow fever virus transmission potential. Science (80-.). 361, 894–899 (2018).

37. J. Quick, N. J. Loman, S. Duraffour, J. T. Simpson, E. Severi, L. Cowley, J. A. Bore, R. Koundouno, G. Dudas, A. Mikhail, N. Ouédraogo, B. Afrough, A. Bah, J. H. J. Baum, B. Becker-Ziaja, J. P. Boettcher, M. Cabeza-Cabrerizo, Á. Camino-Sánchez, L. L. Carter, J. Doerrbecker, T. Enkirch, I. G.-Dorival, N. Hetzelt, J. Hinzmann, T. Holm, L. E. Kafetzopoulou, M. Koropogui, A. Kosgey, E. Kuisma, C. H. Logue, A. Mazzarelli, S. Meisel, M. Mertens, J. Michel, D. Ngabo, K. Nitzsche, E. Pallasch, L. V. Patrono, J. Portmann, J. G. Repits, N. Y. Rickett, A. Sachse, K. Singethan, I. Vitoriano, R. L. Yemanaberhan, E. G. Zekeng, T. Racine, A. Bello, A. A. Sall, O. Faye, O. Faye, N. Magassouba, C. V. Williams, V. Amburgey, L. Winona, E. Davis, J. Gerlach, F. Washington, V. Monteil, M. Jourdain, M. Bererd, A. Camara, H. Somlare, A. Camara, M. Gerard, G. Bado, B. Baillet, D. Delaune, K. Y. Nebie, A. Diarra, Y. Savane, R. B. Pallawo, G. J. Gutierrez, N. Milhano, I. Roger, C. J. Williams, F. Yattara, K. Lewandowski, J. Taylor, P. Rachwal, D. J. Turner, G. Pollakis, J. A. Hiscox, D. A. Matthews, M. K. O. Shea, A. M. Johnston, D. Wilson, E. Hutley, E. Smit, A. Di Caro, R. Wölfel, K. Stoecker, E. Fleischmann, M. Gabriel, S. A. Weller, L. Koivogui, B. Diallo, S. Keïta, A. Rambaut, P. Formenty, S. Günther, M. W. Carroll, Real-time, portable genome sequencing for Ebola surveillance. Nature. 530, 228–232 (2016).

38. R. S. Sikkema, S. D. Pas, D. F. Nieuwenhuijse, Á. O’Toole, J. Verweij, A. van der Linden, Chestakova, C. Schapendonk, M. Pronk, P. Lexmond, T. Bestebroer, R. J. Overmars, S. van Nieuwkoop, W. van den Bijllaardt, R. G. Bentvelsen, M. M. L. van Rijen, A. G. M. Buiting, A. J. G. van Oudheusden, B. M. Diederen, A. M. C. Bergmans, A. van der Eijk, R. Molenkamp, A. Rambaut, A. Timen, J. A. J. W. Kluytmans, B. B. Oude Munnink, M. F. Q. Kluytmans van den Bergh, M. P. G. Koopmans, COVID-19 in health-care workers in three hospitals in the south of the Netherlands: a cross-sectional study. Lancet Infect. Dis. 0 (2020), doi: 10.1016/S1473-3099(20)30527-2.

39. RIVM, Signaleringsoverleg zoönosen | RIVM, (available at https://www.rivm.nl/surveillance-van-infectieziekten/signalering-infectieziekten/signaleringsoverleg-zoonosen).

40. RIVM, Outbreak Management Team (OMT) | RIVM, (available at https://www.rivm.nl/coronavirus-covid-19/omt).

41. A. Kroneman, H. Vennema, K. Deforche, H. v d Avoort, S. Penaranda, M. S. Oberste, J. Vinje, M. Koopmans, An automated genotyping tool for enteroviruses and noroviruses. J Clin Virol. 51, 121–125 (2011).

42. V. M. Corman, O. Landt, M. Kaiser, R. Molenkamp, A. Meijer, D. K. Chu, T. Bleicker, S. Brünink, J. Schneider, M. L. Schmidt, D. G. Mulders, B. L. Haagmans, B. van der Veer, S. van den Brink, L. Wijsman, G. Goderski, J.-L. Romette, J. Ellis, M. Zambon, M. Peiris, H. Goossens, C. Reusken, M. P. Koopmans, C. Drosten, Detection of 2019 novel coronavirus (2019-nCoV) by real-time RT-PCR. Eurosurveillance. 25, 2000045 (2020).

43. C. H. GeurtsvanKessel, N. M. A. Okba, Z. Igloi, S. Bogers, C. W. E. Embregts, B. M. Laksono, L. Leijten, C. Rokx, B. Rijnders, J. Rahamat-Langendoen, J. P. C. van den Akker, J. J. A. van Kampen, A. A. van der Eijk, R. S. van Binnendijk, B. Haagmans, M. Koopmans, An evaluation of COVID-19 serological assays informs future diagnostics and exposure assessment. Nat. Commun. 11, 1–5 (2020).

44. B. B. Oude Munnink, D. F. Nieuwenhuijse, R. S. Sikkema, M. Koopmans, Validating Whole Genome Nanopore Sequencing, using Usutu Virus as an Example. J. Vis. Exp., e60906 (2020).

45. R. C. Edgar, MUSCLE: multiple sequence alignment with high accuracy and high throughput. Nucleic Acids Res. 32, 1792–1797 (2004).

46. L. T. Nguyen, H. A. Schmidt, A. von Haeseler, B. Q. Minh, IQ-TREE: a fast and effective stochastic algorithm for estimating maximum-likelihood phylogenies. Mol Biol Evol. 32, 268–274 (2015).

47. Y. Shu, J. McCauley, GISAID: Global initiative on sharing all influenza data – from vision to reality. Eurosurveillance. 22 (2017),, doi: 10.2807/1560-7917.ES.2017.22.13.30494.

48. H. Wickham, ggplot2: Elegant Graphics for Data Analysis (Springer-Verlag New York, 2016).

49. C.-C. B. voor de Statistiek, Wijken buurtkaart 2019, (available at https://www.cbs.nl/nl-nl/dossier/nederland-regionaal/geografische-data/wijk-en-buurtkaart-2019).

50. P. E, B. RS, Classes and Methods for Spatial Data: the sp Package. R News, 9–13 (2005).

51. GitHub - rspatial/raster: R raster package, (available at https://github.com/rspatial/raster/).

52. cran/rgdal, (available at https://github.com/cran/rgdal).

53. D. S. Candido, I. M. Claro, J. G. de Jesus, W. M. Souza, F. R. R. Moreira, S. Dellicour, T. A. Mellan, L. du Plessis, R. H. M. Pereira, F. C. S. Sales, E. R. Manuli, J. Thézé, L. Almeida, M. T. Menezes, C. M. Voloch, M. J. Fumagalli, T. M. Coletti, C. A. M. da Silva, M. S. Ramundo, M. R. Amorim, H. H. Hoeltgebaum, S. Mishra, M. S. Gill, L. M. Carvalho, L. F. Buss, C. A. Prete, J. Ashworth, H. I. Nakaya, P. S. Peixoto, O. J. Brady, S. M. Nicholls, A. Tanuri, Á. D. Rossi, C. K. V. Braga, A. L. Gerber, A. P. de C. Guimarães, N. Gaburo, C. S. Alencar, A. C. S. Ferreira, C. X. Lima, J. E. Levi, C. Granato, G. M. Ferreira, R. S. Francisco, F. Granja, M. T. Garcia, M. L. Moretti, M. W. Perroud, T. M. P. P. Castiñeiras, C. S. Lazari, S. C. Hill, A. A. de Souza Santos, C. L. Simeoni, J. Forato, A. C. Sposito, A. Z. Schreiber, M. N. N. Santos, C. Z. de Sá, R. P. Souza, L. C. Resende-Moreira, M. M. Teixeira, J. Hubner, P. A. F. Leme, R. G. Moreira, M. L. Nogueira, N. M. Ferguson, S. F. Costa, J. L. Proenca-Modena, A. T. R. Vasconcelos, S. Bhatt, P. Lemey, C.-H. Wu, A. Rambaut, N. J. Loman, R. S. Aguiar, O. G. Pybus, E. C. Sabino, N. R. Faria, Evolution and epidemic spread of SARS-CoV-2 in Brazil. Science (80-.)., eabd2161 (2020).

54. T. Ganyani, C. Kremer, D. Chen, A. Torneri, C. Faes, J. Wallinga, N. Hens, Estimating the generation interval for coronavirus disease (COVID-19) based on symptom onset data, March 2020. Eurosurveillance. 25, 2000257 (2020).

55. B. Korber, W. M. Fischer, S. Gnanakaran, H. Yoon, J. Theiler, W. Abfalterer, N. Hengartner, E. E. Giorgi, T. Bhattacharya, B. Foley, K. M. Hastie, M. D. Parker, D. G. Partridge, C. M. Evans, T. M. Freeman, T. I. de Silva, A. Angyal, R. L. Brown, L. Carrilero, L. R. Green, D. C. Groves, K. J. Johnson, A. J. Keeley, B. B. Lindsey, P. J. Parsons, M. Raza, S. Rowland-Jones, N. Smith, R. M. Tucker, D. Wang, M. D. Wyles, C. McDanal, L. G. Perez, H. Tang, A. Moon-Walker, S. P. Whelan, C. C. LaBranche, E. O. Saphire, D. C. Montefiori, Tracking Changes in SARS-CoV-2 Spike: Evidence that D614G Increases Infectivity of the COVID-19 Virus. Cell. 182, 812-827.e19 (2020).

56. E. M. Volz, medRxiv, in press, doi: 10.1101/2020.07.31.20166082.

57. Promed Post – ProMED-mail, (available at https://promedmail.org/promed-post/?id=20200617.7479510).

58. Promed Post – ProMED-mail, (available at https://promedmail.org/promed-post/?id=7584560).

59. E. Cahan, COVID-19 hits U.S. mink farms after ripping through Europe. Science (80-.). (2020), doi: 10.1126/science.abe3870.

60. (No Title), (available at https://www.actasia.org/wp-content/uploads/2019/10/China-Fur-Report-7.4-DIGITAL-2.pdf).

